# Little skate genome exposes the gene regulatory mechanisms underlying the evolution of vertebrate locomotion

**DOI:** 10.1101/2022.03.14.484236

**Authors:** DongAhn Yoo, Chul Lee, JunHee Park, Young Ho Lee, Adriana Heguy, Jeremy S. Dasen, Heebal Kim, Myungin Baek

## Abstract

The little skate *Leucoraja erinacea*, a cartilaginous fish, displays pelvic fin driven walking-like behaviors using genetic programs and neuronal subtypes similar to those of land vertebrates. However, mechanistic studies on little skate motor circuit development have been limited, due to a lack of high-quality reference genome. Here, we generated an assembly of the little skate genome, containing precise gene annotation and structures, which allowed post-genome analysis of spinal motor neurons (MNs) essential for locomotion. Through interspecies comparison of mouse, skate and chicken MN transcriptomes, shared and divergent MN expression profiles were identified. Conserved MN genes were enriched for early-stage nervous system development. Comparison of accessible chromatin regions between mouse and skate MNs revealed conservation of the potential regulators with divergent transcription factor (TF) networks through which expression of MN genes is differentially regulated. TF networks in little skate MNs are much simpler than those in mouse MNs, suggesting a more fine-grained control of gene expression operates in mouse MNs. These findings suggest conserved and divergent mechanisms controlling MN development system of vertebrates during evolution and the contribution of intricate gene regulatory networks in the emergence of sophisticated motor system in tetrapods.

## Introduction

The little skate *Leucoraja erinacea* is a marine species of jawed vertebrate which shared a common ancestor with tetrapods 473 MYA^1^. Yet, it displays a walking-like behavior which resembles limb-based locomotion^2–4^, suggesting that the genetic programs and neural subtypes essential for walking existed in a common ancestor of cartilaginous fish and tetrapods. Hence, *Leucoraja erinacea* can provide a useful model organism for the study of locomotor circuits, as it may reveal insights into the genetic programs underlying limb-driven locomotion, that may have originated from the common ancestor of vertebrates with paired appendages^5^. Previous developmental studies on the little skate demonstrated a conserved *Hox* TF-dependent regulatory network that specifies MNs innervating fin and limb muscle^4^. Little skate pectoral and pelvic fin MNs express forelimb MN *Hox* genes, *Hoxa6/7* and a hindlimb MN *Hox* gene, *Hoxa10*, respectively. In addition, the little skate genome lacks the *HoxC* cluster^6^ and its spinal cord is occupied by fin MNs without an inter-fin region, which is reminiscent of the *HoxC* cluster mutant mouse^7^. However, the mechanisms that regulate expression of genes controlling motor circuit development remain to be identified, and how different species generate MN subtypes to innervate ∼10 pelvic fin muscles^8^ versus ∼50 limb muscles^9–11^ remains to be determined.

To investigate common and divergent regulatory gene networks of the little skate and tetrapods, integrated analysis of comparative genomics, transcriptomics and epigenomics analyses, including assay for transposase-accessible chromatin using sequencing (ATAC-seq) can be applied. However, in little skate, such genome-wide omics studies have been limited due to severe fragmentation of the reference genome^12^. Limitations of short-read sequencing, such as the 500bp read length in the case of skate genome, are unable to cover long repeats, leading to poor assembly of repeat-rich regions. Continuous long read (CLR) sequencing provides a solution to such assembly errors. With the assistance of available high-quality reference assemblies of phylogenetically close species long read sequencing allows for reference-guided scaffolding and comparative annotations approaches^14, 15^.

Here, we generated a new genome assembly of 2.13 Gb and gene annotation of little skate by applying a combination of PacBio long-read and Illumina short-read sequencing, a state-of-art assembly and annotation pipeline^14–16^. Based on the new reference genome, transcriptome and ATAC-seq analyses were performed to identify gene expression patterns and chromatin accessibility of fin-innervating MNs. Through a comparative transcriptome analysis with two well-studied tetrapods, we characterized both conserved and divergent gene expression patterns, finding that genes controlling neurodevelopment are shared in MNs of different species. Comparative chromatin accessibility analysis revealed that TF regulatory networks in mouse MNs are more complicated than the ones in little skate MNs, which suggest an underlying mechanism of the emergence of intricate motor systems. The findings of this study provide deeper insights into the origin of gene regulatory mechanisms underlying limb-based locomotion.

## Results

### High-quality little skate genome assembly through a combination of long and short read sequencing

The genome of *Leucoraja erinacea* was assembled using a combination of PacBio long read and Illumina short read data (Fig. 1A-B). Presented in Fig. 1A are the top 49 scaffolds ranked by the length which is equivalent to the number of chromosomes in thorny skate genome (sAmbRad1)^13^. The 49 scaffolds constituted 97.8% of the little skate genome. Compared to the previous assembly, our assembled genome size increased by 36.5% to 2.13 Gb^12^; ∼93% of the estimated genome size based on K-mer spectra of whole genome sequencing data (Fig. S1). The contiguity of the genome was also improved by over 300-fold, in terms of contig N50 of 214 Kb. The completeness of the new genome assembly is highlighted in the benchmarking universal single-copy orthologs (BUSCO) gene contents (Fig. S2-3). Missing and fragmented BUSCO which was 69.1% and 23.9% in the previous assembly has been reduced to 7.3% and 3.6% in our assembly, respectively. Also, the proportion of complete BUSCO genes (89.1%) as well as the long transcript lengths demonstrate the completeness of the new assembly compared to the previous one (Fig. S3-4). As repetitive elements are reported to have large variation among cartilaginous fish genomes^13, 17^, the repeats were investigated using Repeatmasker^18^ and were compared. Approximately, 63.6% of the little skate genome is occupied by repetitive elements (Fig.S3B), which may account for its relatively large genome size compared to elephant shark and zebrafish genomes (C.milii-6.1.3 and GRCz11)^19, 20^. Among the repeats, long interspersed nuclear elements (LINE) were the most abundant of all classes which is similar to thorny skate and three shark genome assemblies^17^.

**Figure 1.**
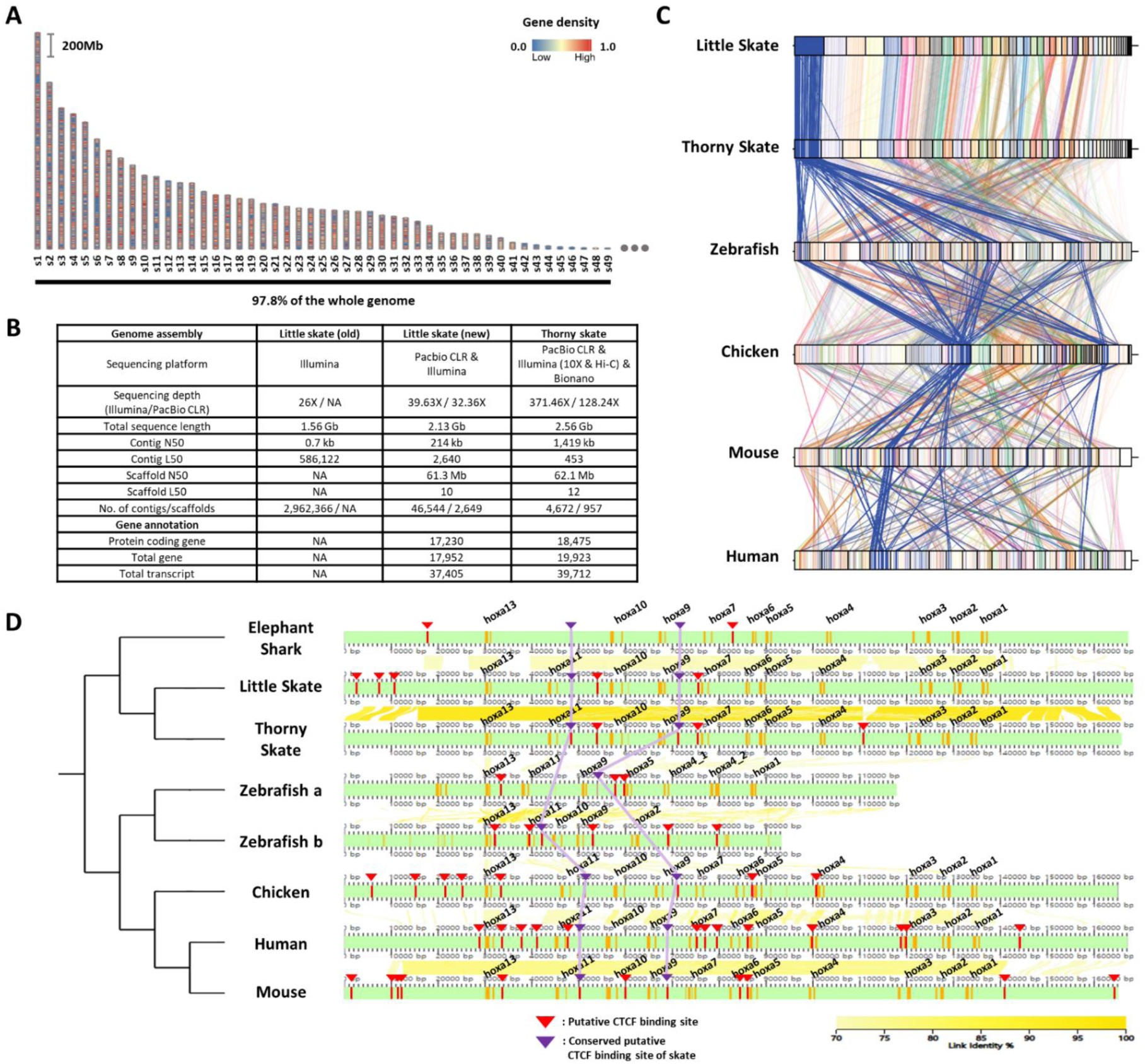
Assembly and comparison of little skate genomes across diverse phylogenetic lineages. A. Top 49 scaffolds ranked by the length. Gene density in the scaffolds is color coded. B. Assembly statistics of the skate genomes. C. Comparison of BUSCO gene synteny across diverse phylogenetic lineages. The BUSCO genes in each scaffold of the new little skate genome are color coded and the longest scaffold (s1) is highlighted in blue. D. Putative binding sites of CTCF in the *HoxA* cluster. Putative CTCF motifs are predicted (described in materials and methods) and indicated by red bars and arrowheads and the ones conserved by purple bars and arrowheads. The exons in the genome is denoted by orange bars. The sequence identities of alignment blocks are indicated by the intensity of yellow color.

A scaffold N50 of 61.3Mb of our little skate genome assembly was comparable to a recently published high-quality thorny skate genome assembly^13^ (62.1Mb; Fig. 1B). The number of protein-coding genes (17,230) and transcripts (17,952) predicted in the new genome assembly was similar to the number in the thorny skate genome (Fig. 1B). As a qualitative assessment of our genome assembly, we compared an existing sequence of the *HoxA* cluster constructed using a BAC clone^21^ (FJ944024), to our assembled genome. Coding sequences in our assembled *HoxA* cluster aligned well with the fully sequenced BAC clone (Fig. S5), highlighting the reliability of the new assembly. We observed complete loss of *HoxC* cluster in the little skate genome which was also shown in the thorny skate genome^6, 13, 22^ and relatively high repeat contents nearby genomic regions in the scaffold41 (s41), which contains the largest number of the *HoxC* neighbor genes (Fig. S6). Although a residual fragment of *HoxC* cluster was reported in the shark genomes^17^, cloudy cat shark, bamboo shark and whale shark genomes are not visualized here due to an absence of gene annotation files.

Comparing the new little skate genome with the one of thorny skate, we found high level of conservation between the species in terms of BUSCO gene contents (Fig. 1C). Interestingly, we also observed that the BUSCO gene synteny of little skate showed more similar patterns with chicken and mammals than with zebrafish (Fig. 1C). In addition to gene contents, we also compared putative CTCF binding sites, which are known to regulate expression of *HoxA* genes during development^23, 24^ (Fig. 1D). In tetrapods, *HoxA* genes are expressed in spinal cord MNs and specify their subtype identities^7, 25, 26^, and similarly in little skate, *HoxA* genes are expressed in spinal cord MN subtypes^4^. Among previously reported CTCF sites in mouse^24^, one located between *Hoxa7* and *Hoxa9*, and between *Hoxa10* and *Hoxa11* were conserved across diverse species and were also observed in little skate *HoxA* cluster despite the phylogenetic distance (Fig. 1D). On the other hand, the CTCF binding sites near *Hoxa13* upstream and between *Hoxa5* and *Hoxa6* which are conserved in other species were not found in putative CTCF sites of skate, elephant shark, and other sharks reported previously^17^. The absence of the CTCF site between *Hoxa5* and *Hoxa6* may contribute to the relatively expanded domain of *Hoxa9* in fin innervating MNs^4, 24^.

### DEG analysis with a new reference genome revealed comprehensive MN markers

Previous RNA sequencing data of little skate MNs^4^, which was analyzed using the zebrafish transcriptome, was re-analyzed with the new little skate genome. Comparing the expression of 10,270 genes in pectoral fin MNs (pec-MNs) with tail-region spinal cord (tail-SC) identified 411 differentially expressed genes (DEGs) including 135 genes upregulated in the pec-MNs and 276 genes in the tail-SC (Fig. 2). Among the identified DEGs, genes associated with the GO term “DNA-binding transcription factor activity” including *Hox10-13* genes were highly expressed in tail-SC compared to pec-MNs while *Hoxa7* was highly expressed in pec-MNs (Fig. 2C), which is consistent with previous immunohistochemical analyses^4^.

**Figure 2.**
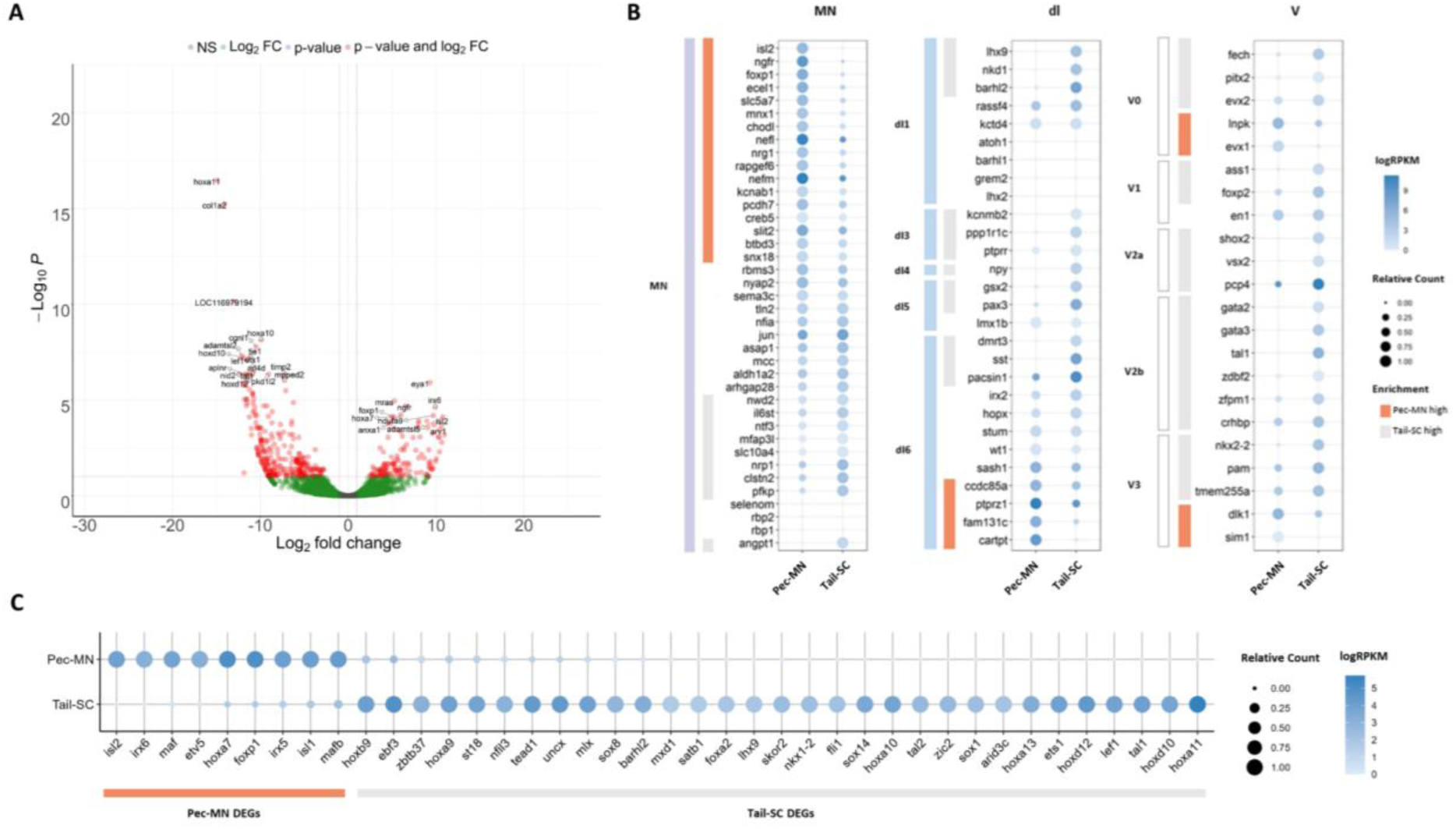
Differentially expressed genes of Pec-MNs and Tail-SC. A. Volcano plot of gene expression. Each dot represents individual genes and the highly expressed genes (FC≥2, adjusted p-value < 0.1) are indicated in red and the top 20 genes with lowest p-value are labelled. The genes highly expressed in Pec-MNs and tail-SC are on the right and left, respectively. B. Comparison of Pec-MN and tail-SC DEGs with the gene expression of mouse MN and interneuron marker genes^27^; MN: Motor neuron, dl: dorsal interneurons (dorsal interneuron subtypes are labeled on the left of light blue bars), V: Ventral interneurons (ventral interneuron subtypes are labeled on the left of white bars). MN and interneuron markers highly expressed in Pec-MNs compared to the tail-SC are indicated by orange bars; while markers expressed at higher levels in the tail-SC than in Pec-MNs are indicated by grey bars. Absolute expression levels are coded in blue and relatively expression levels are indicated by circle size. C. The differentially expressed TFs under the GO term “DNA-binding transcription factor activity”. Pec-MN and tail-SC DEGs are indicated by orange and grey bars, respectively.

Using our newly annotated genome, we comprehensively examined the similarity of gene expression patterns between skate and tetrapod spinal neurons. A set of molecular markers for spinal MNs and interneurons of mouse embryos^27^ were compared with the ones of skate pec-MNs and tail-SC. Overall, we observed a greater number of mouse MN genes highly expressed in pec-MNs (pec-MN vs. tail-SC fold change (FC) ≥ 2; 17 genes) compared with tail-SC (FC ≥ 2; 9 genes; Fig. 2B). On the other hand, a greater number of interneuron markers was observed in tail-SC (FC ≥ 2; 29 genes), which is composed predominantly of interneuron cell-types, than in pec-MNs (FC ≥ 2; 8 genes).

The evolution of genetic programs in MNs was tested by unbiasedly comparing highly expressed genes in pec-MNs of skate with the ones from MNs of mouse and chicken, two well-studied tetrapod species. Genes highly expressed by MNs in mouse were identified by comparing the previously reported transcriptome of mouse MNs at embryonic day I 13.5 to sensory neurons (SNs) at e14.5^28^. Chick embryonic transcriptome data was generated, and the genes highly expressed in chick MNs were identified by comparing the transcriptome of brachial MNs (br-MNs) with that of brachial spinal cord at HH25^29^. Comparing the DEGs, MN genes specifically lost in chick and skate were investigated using the mouse genome as the reference. Several MN genes were specifically lost in skate, while no MN genes were found to be lost in chick (Table. S1). Moreover, comparison of orthologous MN DEGs revealed shared and species-specific expression (Fig. 3A). Among the shared DEGs, 40, 23, and 62 genes were identified in the skate and mouse pair, skate-chick pair and mouse-chick pair, respectively. The 5 genes that were commonly found in all mouse, skate and chick MN DEGs include *Foxp1, Ret, Slc5a7, Dlgap2* and *Cacng2*. Interestingly, the majority of the MN DEGs was found in a species-specific manner (Fig. 3A); 191, 1,734 and 413 DEGs were exclusively found in skate, mouse and chick MNs, respectively. Gene ontology (GO) analysis revealed that neurodevelopment-related terms constituted large proportions among the shared DEGs (Fig. 3B). Among the species-specific MN DEGs, the GO terms associated with neurodevelopment were also identified along with many terms related to cell morphogenesis and transport. Species-specific DEGs in mouse MNs and chick br-MNs include more terms related to neurodevelopment than skate-specific MN DEGs (Fig. 3C).

**Figure 3.**
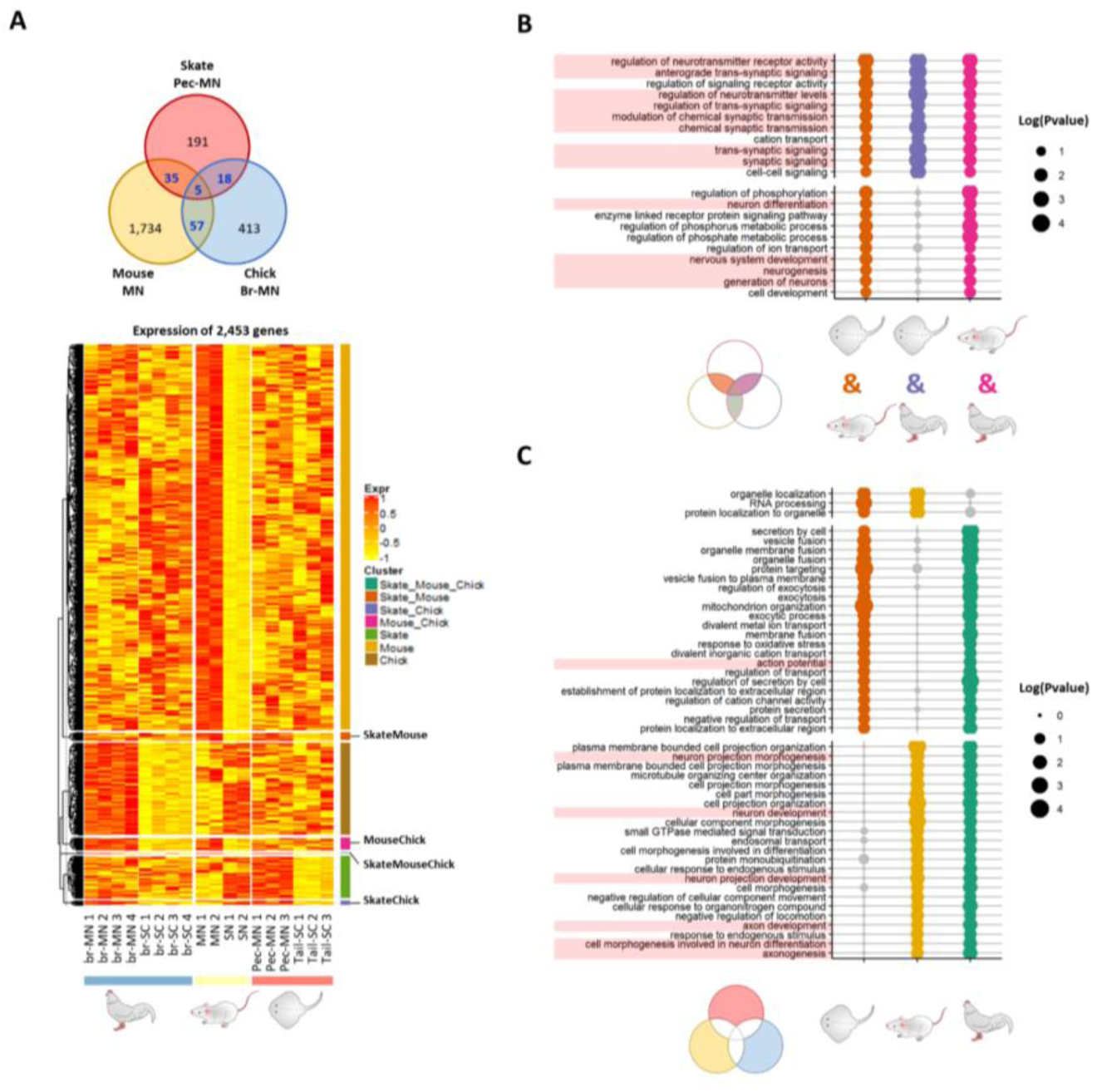
Potential MN marker genes commonly or divergently expressed in different species. A. The Venn diagram and heatmap of MN DEGs of the three species showing shared and species-specific DEGs of little skate, mouse and chick. In the heatmap, expression levels are color coded according to the z-scores in each species. B, C. Biological process GO term enrichment analyses. GO terms identified in more than two species are shown (total list in Additional Data 3. GO term enrichment). Terms related to neurons are highlighted in red on the left. Significance is indicated by circle size. Insignificant GO terms (p≥0.05) are indicated by grey color. B. Enriched GO terms in shared MN DEGs. C. Enriched GO terms in divergent MN DEGs.

### ATAC-seq analysis in little skate MNs uncovered putative regulators of MN DEGs

Our high-quality annotated little skate genome assembly, with sufficient contiguity of scaffold covering potential regulatory regions, allowed us to further investigate the mechanisms of gene regulation in little skate fin MNs. We examined chromatin accessibility by performing ATAC-seq from isolated pectoral/pelvic fin MNs (fin-MNs) and tail-SC. The quality control for ATAC-seq data is summarized in Fig. S7; the highest ATAC-seq read depth was observed right before the transcription start site (TSS) for nucleosome-free reads than the nucleosome-bound reads (Fig. S7A-D). The distribution of insert size shows similar trends with the typical size distribution of ATAC-seq analysis^30^ (Fig. S7E-F). Through peak calling, a total of 61,997 consensus accessible chromatin regions (ACRs) were identified in fin-MNs and tail-SC, indicated by intense red color in the heatmap (Fig. 4A). Among them, 7,869 ACRs were found in both fin-MNs and tail-SC while 35,741 and 18,387 ACRs were found only in fin-MNs and tail-SC, respectively (Fig. 4A and Fig. S8). On average, higher read depth was observed for fin-MN ACRs than that of the tail-SC.

**Figure 4.**
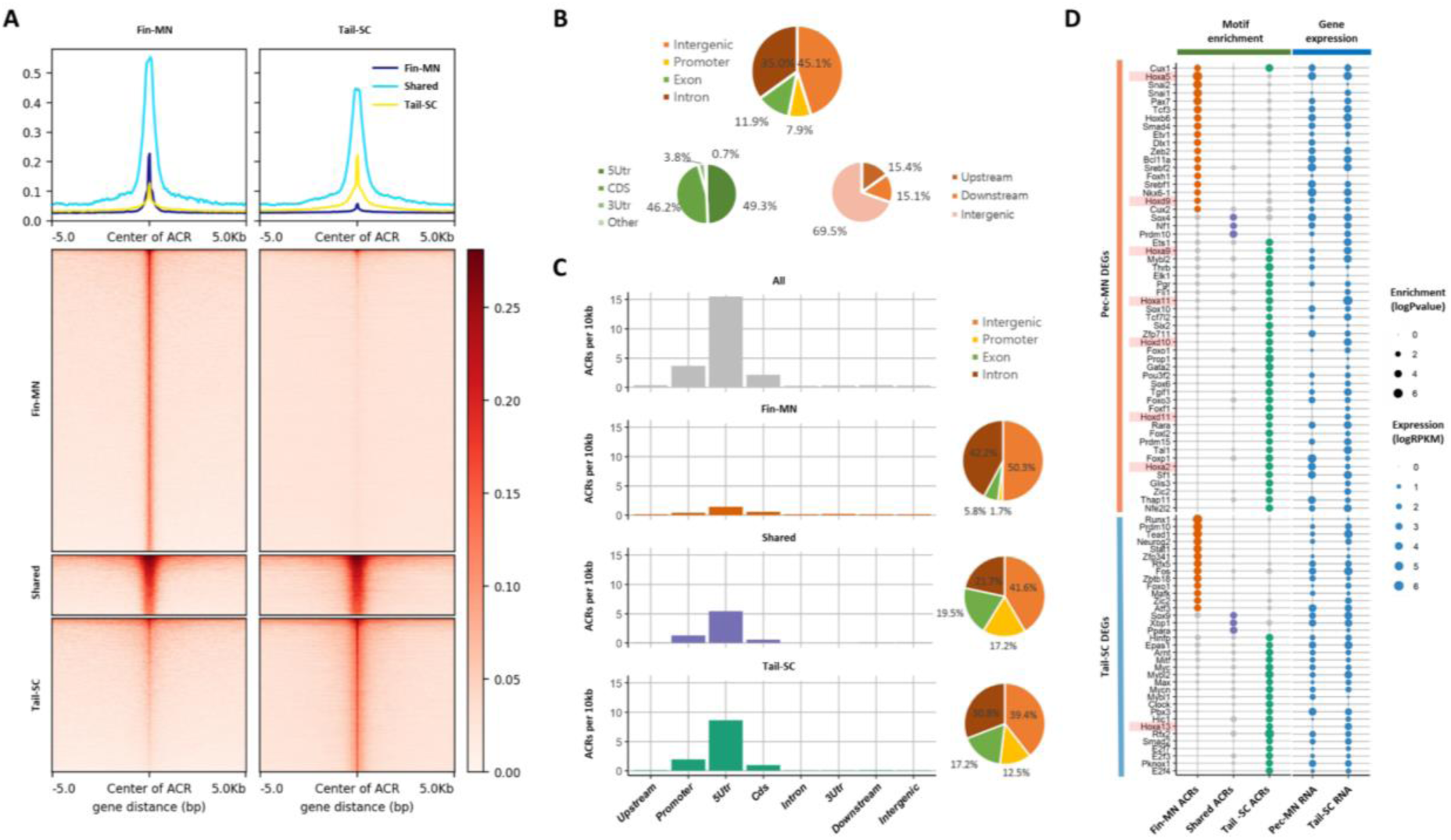
Overview of ATAC-seq data. A. Heatmap of 61997 ACRs. Fin-MN ACRs on the left and tail-SC ACRs on the right. The number of ACRs: both fin-MN and tail-SC, 7869; fin-MNs specific, 35741; tail-SC specific, 18387. Higher intensity of red shows regions with higher ATAC-seq read depth. The line plot on top shows the average read depth around 5 Kb of ACR centers. B. Proportion of all ACRs (fin-MNs and tail-SC) in different genomic regions. C. The density of ACRs (all, fin-MN specific, shared, and tail-SC specific ACRs) in each genomic region normalized by the respective length. D. Significantly enriched TF binding motifs in ACRs (p < 0.05; around 10 Kb up-and downstream of transcription start and end sites of each gene) of DEGs (pec-MN DEGs, upper rows; tail-SC DEGs, lower rows) identified in Fig. 2. From left to right columns: fin-MN specific, shared and tail-SC specific ACRs. The respective expression in pec-MNs and tail-SC is also shown for each TF on the right two columns. Significance and expression levels are indicated by circle size.

The genome-wide distribution analysis of ACRs showed that many ACRs were found in intron and intergenic regions for all ACR types (intron, 35.0%; intergenic region, 45.1%; Fig. 4B); however, normalizing each region by its length, the density of ACRs was found to be highest in promoters and 5’UTRs (Fig. 4C). Even though higher number of fin-MN-specific ACRs were identified, the ACR normalized by region length showed lower density for fin-MN specific ACRs because large number of the ACRs in the fin-MNs were found in intron and intergenic regions which constitute the largest proportion of the genome. After observing that the ACR density is higher near the TSS, the correlation between chromatin accessibility in the promoter region and the gene expression level was investigated. As a result, we found that the genes with greater depth of ATAC-seq in their promoter region are generally expressed at higher levels than those genes with closed chromatin form in both fin-MNs and tail-SC (Fig. S9), indicating that ACRs in the promoter region are likely to be associated with gene activation.

Using the ACR data, enrichment test of putative TF binding sites was performed for pec-MNs and tail-SC DEGs to reveal potential regulatory mechanisms (Fig. 4D). Overall, 18, 3 and 35 TFs were significantly enriched in fin-MN specific ACRs, shared and tail-SC specific ACRs in pec-MN DEGs, respectively. Interestingly, differential enrichment of binding sites of *Hox* proteins, well-known MN subtype regulators along the rostro-caudal axis of the spinal cord^4, 25, 31^, was found in the ACRs of pec-MNs and tail-SC DEGs. Fin-MN-specific ACRs in the pec-MN DEGs were enriched with binding sites of *Hoxa5* and *Hoxd9*, which are expressed in the fin-MNs of little skate^4^. Together with the binding sites of the *Hox* proteins, the binding sites of *Nkx6-1*, a TF regulating progenitor differentiation and MN specification^32, 33^, were enriched in Fin-MN specific ACRs of pec-MN DEGs. On the other hand, the tail-SC-specific ACRs in the pec-MN DEGs were enriched with the binding sites of *Hoxa9*, *Hoxa11*, *Hoxd10* and *Hoxd11*, most of which are expressed in fin-MNs of little skate^4^. Moreover, among the tail-SC DEGs, binding sites of 13, 3 and 19 TFs were significantly enriched in fin-MN-specific, common and tail-SC-specific ACRs. Among them, binding sites of *Neurog2*, a TF essential for MN specification^34^, were enriched in fin MN ACRs while the binding sites of *Hoxa13*, a caudal Hox protein^35^ were enriched in tail-SC-specific ACRs.

### Little skate and mouse hold shared and diverged gene regulatory system in MNs

To evaluate the degree of conservation of the skate gene regulatory system in MNs with that of land-walking vertebrates, the TF binding motifs were investigated for the ACRs in the skate pec-MN DEGs, mouse MN DEGs and the shared DEGs (Fig. 5A, D-E). As a result, binding sites of 16, 20 and 1 TFs were commonly enriched in both mouse and skate MN ACRs of mouse MN DEGs, skate pec-MN DEGs and the shared DEGs, respectively (Fig. 5B). Among the putative binding sites enriched in both mouse and skate MN ACRs, half were insignificantly enriched in mouse SN DEGs. On the other hand, enrichment of these TFs was similar to that of the tail-SC ACRs. In the MN ACRs of MN DEGs, binding sites of 7 TFs—*E2f3, Klf1, Klf3 Klf6 Sp1, Sp2* and *Sp5*—were significantly enriched in both skate and mouse MN DEGs as well as the binding sites of *Snai1* enriched in skate DEGs and shared DEGs, suggesting the possibility of conserved common regulator systems for skate and mouse MN genes.

**Figure 5.**
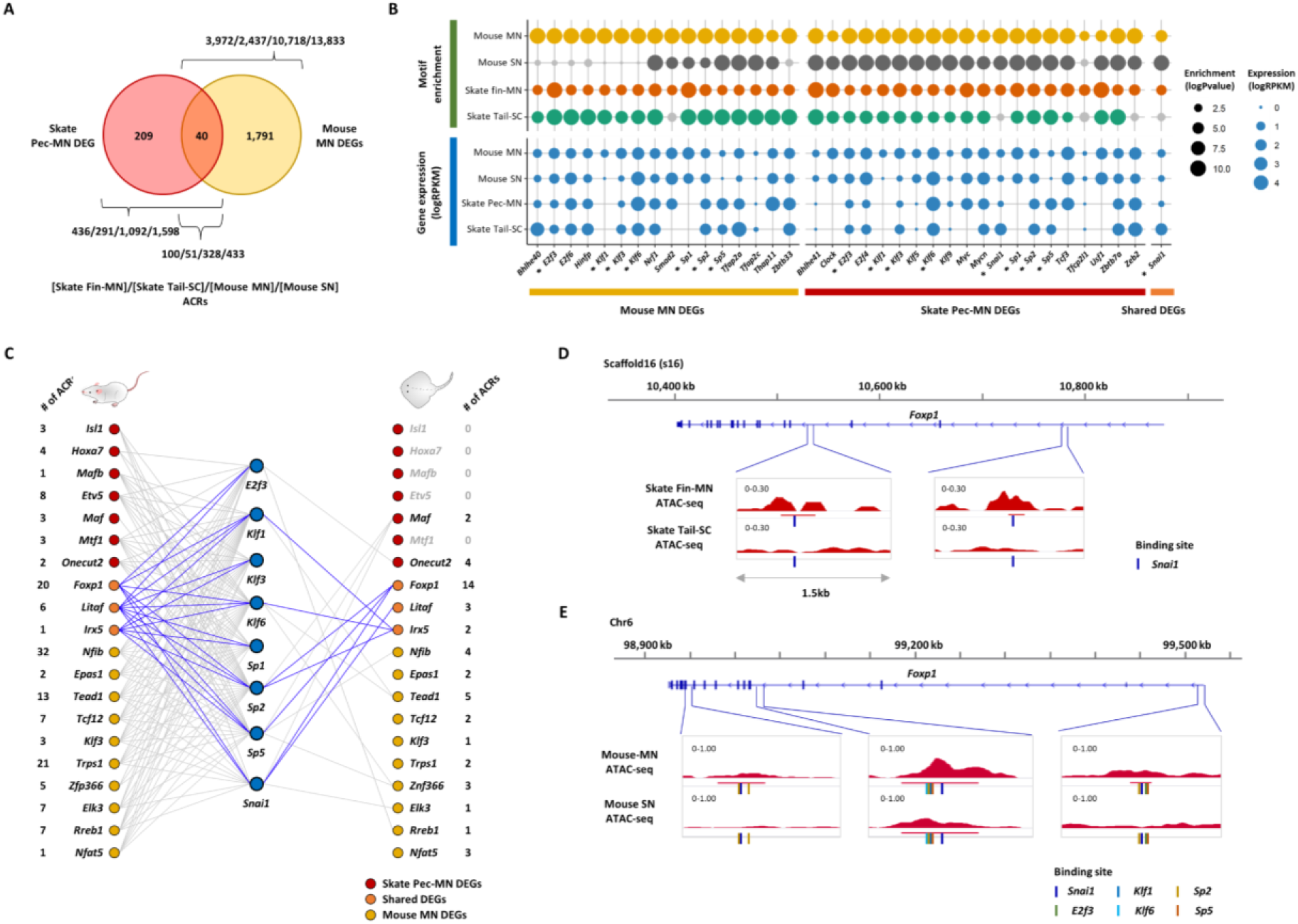
Gene regulatory modules in MNs. A. Number of MN DEGs and ACRs (around 10 Kb up- and downstream of transcription start and end sites of each gene) in MN DEGs of skate and mouse. B. Significantly enriched TF binding motifs (p < 0.05) identified in both skate and mouse MN ACRs are color coded. Non-significant TF binding motifs are denoted by grey. The enriched TF binding sites among skate, mouse or shared MN DEGs are indicated at the bottom. The gene expression of each TF is also shown in lower rows. The enrichment scores and expression levels are indicated by circle size. *Commonly identified TF motifs between different DEGs. C. The potential interactions between the common TFs found in MN ACRs (B) with the top 10 MN-DEG TFs of mouse and skate based on binding sites predicted on ACRs. The putative binding of common TFs on the ACRs of shared MN DEGs are indicated by blue. Number of ACRs in each gene is indicated on the left (mouse) and on the right (skate). Skate MN DEG TFs, red; shared MN DEG TFs, orange; mouse MN DEG TFs, yellow. D, E. The examples of *Snai1* and other binding sites identified in *Foxp1*. Binding motifs for each TF are indicated by color bars. D. Skate *Foxp1*. E. Mouse *Foxp1*.

To investigate the regulatory circuits of the candidate TFs, we examined the top 20 genes of skate pec-MN and mouse MN DEGs (top 10 each, ranked by the p-value) belonging to the GO term “DNA-binding TF activity”. Interaction between the commonly enriched 8 TFs in Fig. 5B with the DEG TFs were predicted by examining the binding sites of the 8 TFs in the ACRs of MN DEG TFs. As a result, highly different TF network was observed between mouse and skate (Fig. 5C-E). The interactions between the TFs also showed highly different profiles for the three shared DEGs as well, including *Foxp1, Litaf* and *Irx5* (Fig. 5C). The example of ACRs with putative *Snai1* binding sites in mouse (Fig. 5E) having a greater number of binding sites than in skate, may indicate more complicated interactions between these TFs in mouse compared to those in skate (Fig. 5D).

## Discussion

The little skate is an emerging model to study the evolution of the neuronal circuits for locomotion. Here, in an attempt to understand the origin of the genetic program for tetrapods locomotion, we generated a new reference genome assembly of little skate with gene annotations. We investigated the regulatory mechanisms of MN gene expression through integrating gene expression and chromatin accessibility data from multiple vertebrate species. As a result, we found both conserved and divergent molecular markers for MNs across multiple species. Moreover, through chromatin accessibility analysis we provide a list of TFs that could regulate the expression of MN genes in mouse and skate. These findings provide deeper insights into the emergence and evolution of gene regulatory pathways necessary for development of the circuits responsible for limb-based locomotion.

### High-quality genome assembly of little skate provides a reference for studying gene regulatory mechanisms in MNs

A previous highly fragmented and incomplete genome of *Leucoraja erinacea*^12^ is unsuitable for post-genome analysis. Therefore, improving the quality of the little skate reference genome is a first step towards a more complete analysis of gene regulatory networks. To improve the quality of the little skate genome, whole genome sequencing data including PacBio long reads and Illumina short reads were generated. Even though this study used fewer resources relative to that of more recent sequencing pipelines, which use multiple sequencing technologies including chromosomal conformation capture and physical maps^13^, this study could assemble a reference genome which is highly contiguous and offers reliable coding and regulatory region data. This was accomplished by using the assembly of a closely-related species—thorny skate—as a reference during scaffolding process. However, it is important to note that the contig N50 of this little skate genome is still lower than that of the thorny skate assembly. Also, the higher number of scaffolds and the presence of little skate contigs which had not been localized to pseudo-chromosomes of thorny skate may indicate that little skate and thorny skate genomes contain a few structural disagreements.

Despite this limitation, the new reference genome allowed for a more reliable RNA-seq analysis. Through re-analyzing the previously generated RNA-seq data^4^, we found genes which are consistent with the previous immunohistochemical analyses^4^. In addition, we also observed that the expression patterns of MN and interneuron markers in mouse^27^ are highly similar in skate, supporting previous conclusions. A greater number of DEGs were found through re-analysis of RNA-seq data using the new reference genome (Fig. 2), which reflects the limitation of the previous analyses using the zebrafish transcriptome. Aligning the new gene annotation data against the previous *de novo* transcriptome, and investigating the distribution of alignment length, we also found that the previous *de novo* transcriptome contains many fragmented genes (Fig. S4A). The previous transcripts generally cover only a small fraction (<20%) of the transcripts predicted in our genome assembly while the new transcripts cover close to 100% of the previous transcriptome. An example case is illustrated in the *Foxp1* gene, a well-known limb/fin MN fate determinant (Fig. S4B). This improved DEG analysis therefore offers a more complete view of gene expression profiles of MNs in little skate.

### The evolution of gene regulatory modules for MN development

The comparative analyses identified both common and divergent gene expression in MNs. A list of conserved putative TF binding sites of mouse and skate MN genes were identified, suggesting that MN gene expression is under the influence of shared regulatory factors. Investigation of conservation of the regulatory system of skate and mouse revealed the candidate shared regulatory factors may control differential gene expression of skate and mouse MN DEGs through both regulating different downstream TFs and differentially regulating shared downstream TFs (Fig. 5C).

During the evolution of motor behaviors, a more complex nervous system likely emerged to control more sophisticated limb movements, such as hand dexterity. Little skate displays walking-like behavior using only around 10 muscles in the pelvic fin, while mammals can use up to 50 muscles in the limb. How the system evolved to generate a more intricate nervous system remains an open question. One mechanism to achieve this might be to increase the number of regulatory modules that control gene expression. Recently, an increase in the size of intergenic regions in neuronal genes was proposed as one plausible mechanism to generate a more complex nervous system^28^. Nature might have invented additional ways of achieving this. Although neuronal genes in little skate have much longer intergenic regions than non-neuronal genes, the intergenic regions in neuronal genes contain a similar fraction of ACRs as in non-neuronal genes as opposed to mouse with significantly increased neuronal gene ACRs (Fig. S10). While birds display very complex behaviors, its genome contains short intergenic regions^36^. In our study, we propose another way of achieving complex nervous system, through more intricate TF networks. As illustrated in the Fig. 5C, a greater number of shared TFs binds to their downstream TFs in mouse MNs than in skate pec-MNs, allowing an intricate control of gene expression and thus the more complex nervous system of mouse compared to that of skate even though the hypothesis remains to be validated in additional organisms.

In summary, the comparative analysis of the current study discovered evidence of species-specific changes apart from the conservation. Divergent expression of MN genes (Fig. 3A) could allow a greater number of motor pools to be generated in mouse to control complex limb structures; loss of *HoxC* cluster in little skate genome (Fig. S6) might be the underlying cause of the expanded fin MN domain in the spinal cord without an inter-fin region; loss of a subset of CTCF binding motifs in *HoxA* cluster of little skate (Fig. 1D) may have induced the relatively expanded domain of *Hoxa9*. The changes in genome structure and sequence could lead to changes in the expression of MN-specific genes as well as *Hox* genes across the rostro-caudal axis of the spinal cord, affecting the MN organization^37–39^. The changes in the MN organization along the rostro-caudal axis of the spinal cord may affect the connectivity of MNs, which would eventually lead to changes in the pattern of locomotion^40, 41^. Continued effort is needed and complete understanding of the shared and species-specific genetic system would elucidate the origin of molecular and neuronal circuit for tetrapods locomotion.

### Limitations

The findings in this study demonstrates the conservation and divergence in gene expression pattern and gene regulatory system among skate and tetrapod species. However, it is important to note that whether the candidate TFs actually bind to the putative binding sites and whether TFs function as activators or repressors are yet to be validated. As another limitation of the current study, the ACRs analyzed here did not consider long-range regulatory elements. For more accurate characterization of regulatory elements, future studies need a more comprehensive approach including ChIP-seq and chromatin conformation capture assay.

## Materials and Methods

### Sampling and DNA extraction and whole genome sequencing

For genomic DNA extraction from little skates (Marine Biological laboratory; 5 stage and sex unmatched skate embryos), the guts were removed and washed 2 times with 1x PBS. Around 500 µl of G2 buffer (Qiagen) with RNase A (200 µg/ml) was added to the tissue, minced with a razor blade, and transferred to 19 ml G2 buffer in a 50 ml falcon tube. 1 ml of Proteinase K (Qiagen, 19131) was added to the tube and vortexed for 5 seconds, followed by incubation for 2 hr at 50 °C. The samples were vortexed for 10 seconds and loaded onto a genomic DNA column (Qiagen, 10262). Column purification was performed following the manufacturer’s protocol. For PacBio long read sequencing, little skate genomic DNA (6.5–9 µg in 150 µl volume) was transferred to a Covaris g-Tube (Covaris, #520079) and sheared in a tabletop centrifuge (Eppendorf, #5424) at 3,600 rpm for 30 seconds. After shearing, DNA was cleaned with 0.4x ratio of Ampure XP magnetic beads (Beckman coulter). Eluted DNA was processed in a PacBio SMRTbell template prep kit v1.0 (PacBio, #100-222-300) following the manufacture’s protocol. The SMRTbell templates larger than 15 Kb were enriched using a Blue Pippen size-selection system (Sage Science, #BLF7510). Following size selection, the DNA was cleaned and concentrated using Ampure Xp magnetic beads. The sequencing was performed on the PacBio Sequel system. For Illumina short read sequencing, 1 µg of the high-quality skate DNA was put into Kapa-Roche Library prep kit. The sample underwent 1 PCR cycle. 150 bp paired-end sequencing was performed on the Hiseq4000 with v4 chemistry (Illumina, CA).

### Chick MN RNA sequencing

For chick MN RNA sequencing, HH stage 12–13 chick embryos were electroporated with a plasmid that drives nuclear YPF expression under the control of *Hb9* promoter^42^. After 3 day incubation, br-level spinal cords were isolated and dissociated as described^43^. 5000–23000 YFP^+^ MNs were collected using a FACS sorter (BD, FACSAria III) and PicoPure RNA Isolation Kit (Arcturus, KIT0202) was used to extract total RNAs. cDNA was synthesized and amplified using Ovation RNA amplification system v2 (Nugen, 3100-12). The sequencing library was prepared using the TruSeq RNA sample prep Kit (Illumina, CA). The suitable fragments (350–450 bp) were selected as templates for PCR amplification using BluePippin 2% agarose gel cassette (Sage Science, MA). 150 bp paired-end sequencing was performed in the NovaSeq6000 (Illumina, CA).

### Collection of public data

The RNA-seq raw data of little skate pec-MNs and tail-SC used in differential gene expression analysis available under accession number PRJNA414974 from the NCBI SRA were downloaded. The RNA and ATAC-seq raw data of mouse MNs at e13.5 and SNs at e14.5 under the bioproject ID of PRJNA630707 were downloaded.

The human (GRCh38), mouse (GRCm38), chicken (GRCg6a), zebrafish (GRCz11) and elephant shark (C.milii-6.1.3) reference genomes were collected from Ensembl FTP. The thorny skate (sAmbRad1) reference genome was downloaded from NCBI. Lastly, the three shark genomes were collected from the previous study^17^.

### Sampling of tissues and generation of ATAC-seq data

In developmental stage 30^44^ little skates, pectoral and pelvic MNs were labeled by injecting rhodamine-dextran (3K, Invitrogen). After 1–2 day incubation at RT, cell dissociation was performed as described^43^ with some modifications. Spinal cords were isolated and digested with pronase (Roche, 11459643001) for 30 min at RT and dissociated MNs from several labeled spinal cords were collected manually and processed for ATAC sequencing library preparation as described^45^. In brief, 150–426 labeled MNs were collected in 25 µl cold ATAC-seq RSB (10 mM Tris-HCl pH7.4, 10 mM NaCl, and 3 mM MgCl_2_ in Nuclease free water) and centrifuged at 500 rcf for 10 min in a pre-chilled fixed-angle centrifuge. The supernatant was removed and 10 µl of transposition mix (3.3 µl PBS, 1.15 µl Nuclease free water, 5 µl 2x TD buffer, 0.25 µl Tn5 (1/10 diluted: 5 µl 2x TD buffer, 4 µl nuclease free water, 1 µl Tn5), 0.1 µl 1% digitonin) was added and resuspended by pipetting six times. The samples were incubated with shaking at 1000 rpm for 30 min at 37 °C. Tagmented DNA was purified with Qiagen minelute reaction cleanup kit (Qiagen, 28206). PCR mixtures (10 µl Nuclease free water, 2.5 µl 25 µM primer (Ad1), 2.5 µl 25 µM primer (Ad2.X), 25 µl 2x NEB master mix, transposed sample 20 µl) were prepared and primary PCR was performed in the following condition: 72 °C 5 min, 1 cycle; 98 °C 30 sec, 1 cycle; 98 °C 10 sec, 63 °C 30 sec, 72 °C 1 min 5 cycles; hold at 4 °C. PCR mixture (3.76 µl nuclease free water, 0.5 µl 25 µM primers (Ad1.1), 0.5 µl 25 µM primers (Ad2.X), 0.24 µl 25x SybrGold in DMSO, 5 µl 2x NEB master mix, 5 µl pre-Amplified sample) were prepared and quantitative PCR was performed as follows: 98 °C 30 sec, 1 cycle; 98 °C 10 sec, 63 °C 30 sec, 72 °C 1 min 20 cycles; hold at 4 °C. 1/4 max cycles were determined (N) and the primary PCR reaction products underwent N more PCR cycles. For the tail-SC, 240–1000 cells from two embryos were used for ATAC sequencing library preparation. The sequencing was performed on a paired end 50 cycle lane of the Hiseq 4000 using v4 chemistry.

The primers used in the ATAC library preparation were as follows:

AD1.1: AATGATACGGCGACCACCGAGATCTACACTAGATCGCTCGTCGGCAGCGTCAGAT GTG; AD2.1: TAAGGCGACAAGCAGAAGACGGCATACGAGATTCGCCTTAGTCTCGTGGG CTCGGAGATGT; AD2.2: CGTACTAGCAAGCAGAAGACGGCATACGAGATCTAGTACGGT CTCGTGGGCTCGGAGATGT; AD2.3: AGGCAGAACAAGCAGAAGACGGCATACGAGATTT CTGCCTGTCTCGTGGGCTCGGAGATGT; AD2.4: TCCTGAGCCAAGCAGAAGACGGCATA CGAGATGCTCAGGAGTCTCGTGGGCTCGGAGATGT; AD2.5: GGACTCCTCAAGCAGAAG ACGGCATACGAGATAGGAGTCCGTCTCGTGGGCTCGGAGATGT

### Genome assembly of little skate

The initial contig assembly of little skate genome was performed using raw PacBio reads as input via Flye (v. 2.7.1)^16^. Default parameter was used for this initial assembly. Haplotypic duplication of the initial contig was identified and removed based on read depth using purge_dups^46^. Scaffolding was performed on the duplicate-removed contigs via reference-guided approach implemented in RaGOO^14^ using the reference genome of thorny skate or *Amblyraja radiata* available from RefSeq accession GCA_010909765.1. The scaffolded genome of little skate was further polished using Illumina short reads with FreeBayes^47^. The vertebrate gene data (vertebrata_odb10) of Benchmarking Universal Single-Copy Orthologs (BUSCO v. 5. 2. 2) was used to evaluate the final assembly^48^. The synteny of BUSCO gene was visualized via ChrOrthLin<u>k (</u>https://github.com/chulbioinfo/chrorthlink/)^49^. The simplified process of genome assembly is summarized in Fig. S2. Comparison among genome assemblies and with previous genome assembly or libraries was made using MashMap^50^.

For gene annotation, comparative annotation approach was used. Cactus (v. 1.0.0)^51^ alignment of thorny skate and little skate genome was performed. Comparative annotation toolkit (CAT)^15^ was run using the annotation of thorny skate (GCA_010909765.1). Repetitive elements were analyzed using Windowmasker^52^. To investigate different classes of repeats, RepeatModeler v2.0.1^53^ was used to create species-specific repeat library using default parameters. Repeat annotation was performed by RepeatMasker v4.1.2^18^.

### Identification of ortholog clusters

Using the amino acid sequence of the protein-coding genes of skate, mouse (GRCm38), chicken (GRCg6a), orthologous gene clusters were investigated. OrthoVenn2 was used to identify the orthologous gene clusters using the e-value of 0.01 and the default inflation value^54^. Among the identified ortholog gene clusters, single-copy orthologs identified in all three species were used for next multiple-species RNA-seq analysis.

### RNA-seq data processing and differential gene expression analysis

Quality of RNA-seq raw reads were checked with FastQC (v. 0.11.9). Next, the raw data was trimmed for low quality and Illumina truseq adapter sequences using trimmomatic-0.39^55^ with the following options: “ILLUMINACLIP:[AdapterFile]:2:30:10 LEADING:20 TRAILING:20 SLIDINGWINDOW:4:20 MINLEN:50”. The resulting clean reads were aligned to the reconstructed little skate genome with STAR (v. 2.7.5a)^56^ and were quantified using featureCounts^57^. To explore differential expression between the two tissues (pec-MNs and tail-SC), statistical test was performed via DESeq2^58^ with generalized linear model. Genes with FDR-adjusted p-value of <0.1 were considered significant^59^. For the visualization of RNA-seq data, bigwig files were generated using deeptools (v. 3.4.3) with CPM read depth normalization. For the comparison of known markers, the specific markers provided by^27^ was used.

For the comparison of gene expression across multiple species, RNA-seq data generated from pec-MNs of little skate, br-MNs of chick and MNs of mouse were used. The comparison was made using 9,253 orthologous genes present in all species. The orthologous genes were further filtered to obtain the list of genes showing minimum expression level of 20 average read counts for each species. To account for different gene length in multiple species, RPKM normalization was used. P-value of 0.05 and the fold change of 1.5 were used to identify MN DEGs in the multiple species RNA-seq data. To control for MN expression, tail-SC in skate was used to calculate Z-score in skate while mouse SNs and chick br-SC were used for mouse and chick, respectively. The visualization of the Z-score transformed gene expression was done via R “ComplexHeatmap” package. The functional enrichment of the candidate genes was investigated using biological process terms of Panther^60^.

### ATAC-seq data processing and differential accessibility analysis

For chromatin accessibility assay, ATAC-seq data was used. Similar to RNA-seq data, the quality of the raw sequence data was investigated with FastQC (v. 0.11.9) followed by trimming of sequences with low quality and NexteraPE adapter sequences using trimmomatic-0.39^55^ with following options: “ILLUMINACLIP:[AdapterFile]:2:30:10 LEADING:20 TRAILING:20 SLIDINGWINDOW:4:20 MINLEN:30”. The clean reads were aligned to reference genome of little skate with BWA (v. 0.7.17). The quality of the ATAC-seq reads were further checked using R “ATACseqQC” package. After the alignment, duplicated reads were removed with Picard (v. 2.23.2) (http://broadinstitute.github.io/picard/). Moreover, reads with low mapping quality and those reads that were originated from MT genome were removed with samtools^61^ (v. 1.9). Peak calling was performed for each tissue after pooling the replicates of respective tissue (fin-MNs and tail-SC) using MACS2 (v. 2.2.7.1) with the following options: “–keep-dup all –nomodel, -q 0.05– -f BAMPE -g 1,974,810,099”. Consensus ACRs were defined by merging the peaks from the two tissues using bedtools merge (v. 2.29.2). For the visualization of ATAC-seq data, bed files of ACRs and bigwig files generated by deeptools (v. 3.4.3) were used.

The ACRs identified were further classified into different regions based on gene annotation (including upstream, promoter, 5’UTR, CDS, intron, 3’UTR, downstream and intergenic). The region covering 1,000 bp upstream of start of a gene was defined as promoter region and 10,000 bp upstream of promoter region or downstream of 3’ end of a gene was defined as upstream or downstream, respectively. ACRs located in the remaining non-genic regions were considered intergenic ACRs.

### TF binding motif analysis

For the ACRs of interest and promoter regions of the candidate genes, motif enrichment analysis was performed with “annotatePeaks.pl” of HOMER software^62^ using the collection of homer vertebrate TFs and the binding motifs of *Hox* gene families available in HOCOMOCO database v11^63^. The putative CTCF binding site prediction was also performed using HOMER vertebrate CTCF binding motifs. Enrichment of each TF binding of a set of gene was tested by comparing the observed binding motif counts in that set with the empirical distribution of binding motif counts generated from 10,000 sets of randomly selected ACRs. The empirical p-value of <0.05 was considered significant.

### Data availability

The genome sequence of little skate is archived on NCBI BioProject under accession PRJNA747829. The RNA-seq data of chick brachial MN, and ATAC-seq data of little skate are available from the NCBI Gene Expression Omnibus database under accession number GSE180337.

### Code availability

This study does not involve previously unreported computer code or algorithms. The list of software, their versions and options used are described in the methods section.

## Acknowledgments

We thank Orly Wapinski for advice with ATAC sequencing; NYU Langone’s Genome Technology Center for technical supports. This study was supported by the DGIST R&D Program of the Ministry of Science and ICT (2021010004) to M.B. It was also supported by Basic Science Research Program through the National Research Foundation of Korea (NRF) funded by the Ministry of Education (NRF-2019R1I1A2A01041345) to M.B., the Marine Biotechnology Program of the Korea Institute of Marine Science and Technology Promotion (KIMST) funded by the Ministry of Ocean and Fisheries (MOF) (No. 20180430), Republic of Korea to HK.

## Author contribution

D.Y., H.K., J.S.D, and M.B conceptualized the project, designed and performed the experiments, and wrote the paper. C.L. and Y.L. analyzed the genome and transcriptome data. A.H. generated skate sequencing data. J.P. generated chick sequencing data. All authors read the paper.

## Conflict of interest

All authors declare no competing interest.

## Supplementary information

**Figure S1.**
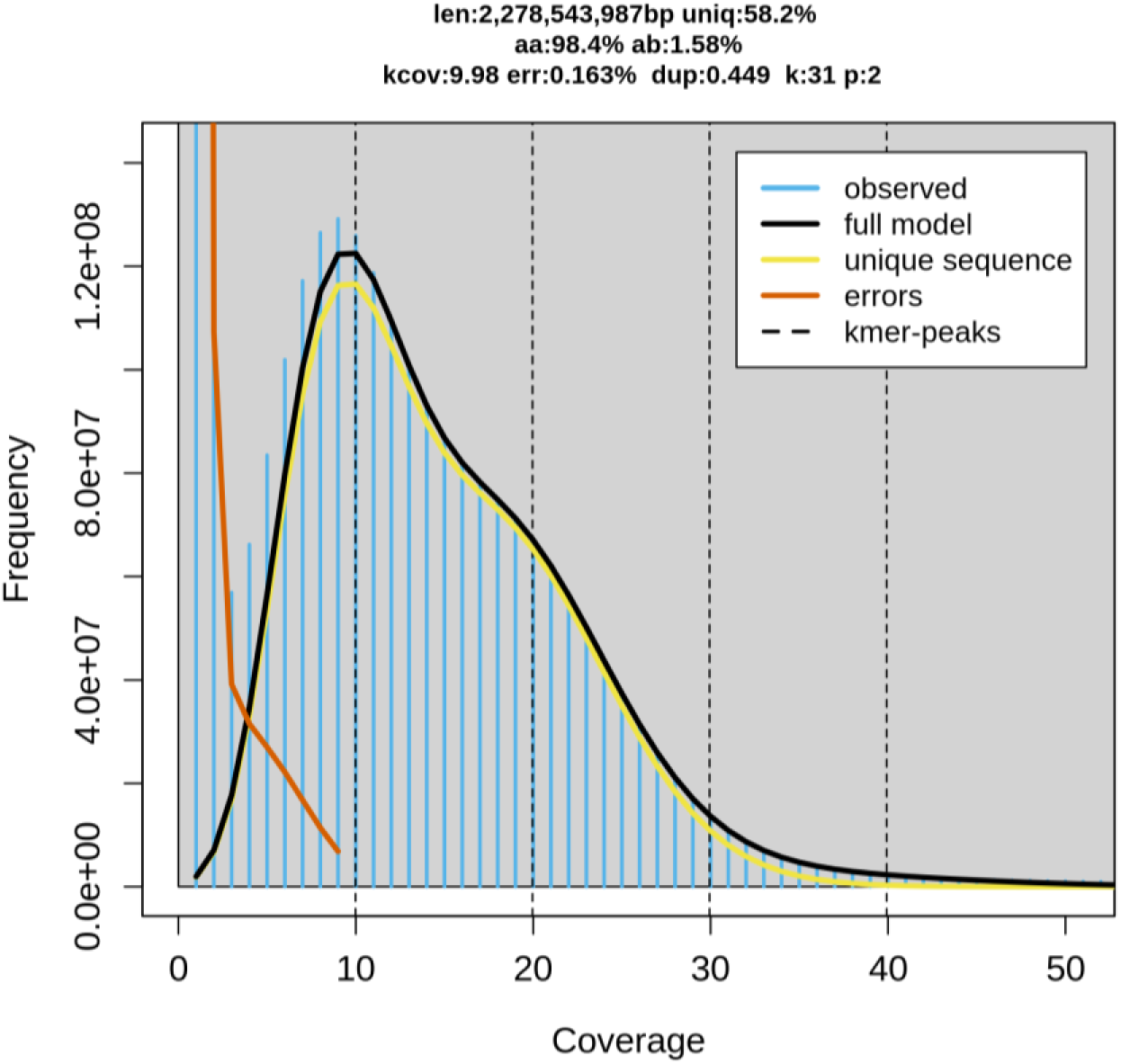
Genome size estimation via Genome Scope 2. K-mer spectrum with K = 31 and the fitted model of little skate genome. (“len”: estimated genome size, “uniq”: proportion of the unique sequence, “aa”: homozygosity, “ab”: heterozygosity, “kcov”: K-mer coverage, “err”: error rate, “dup”: average rate of read duplications.)

**Figure S2.**
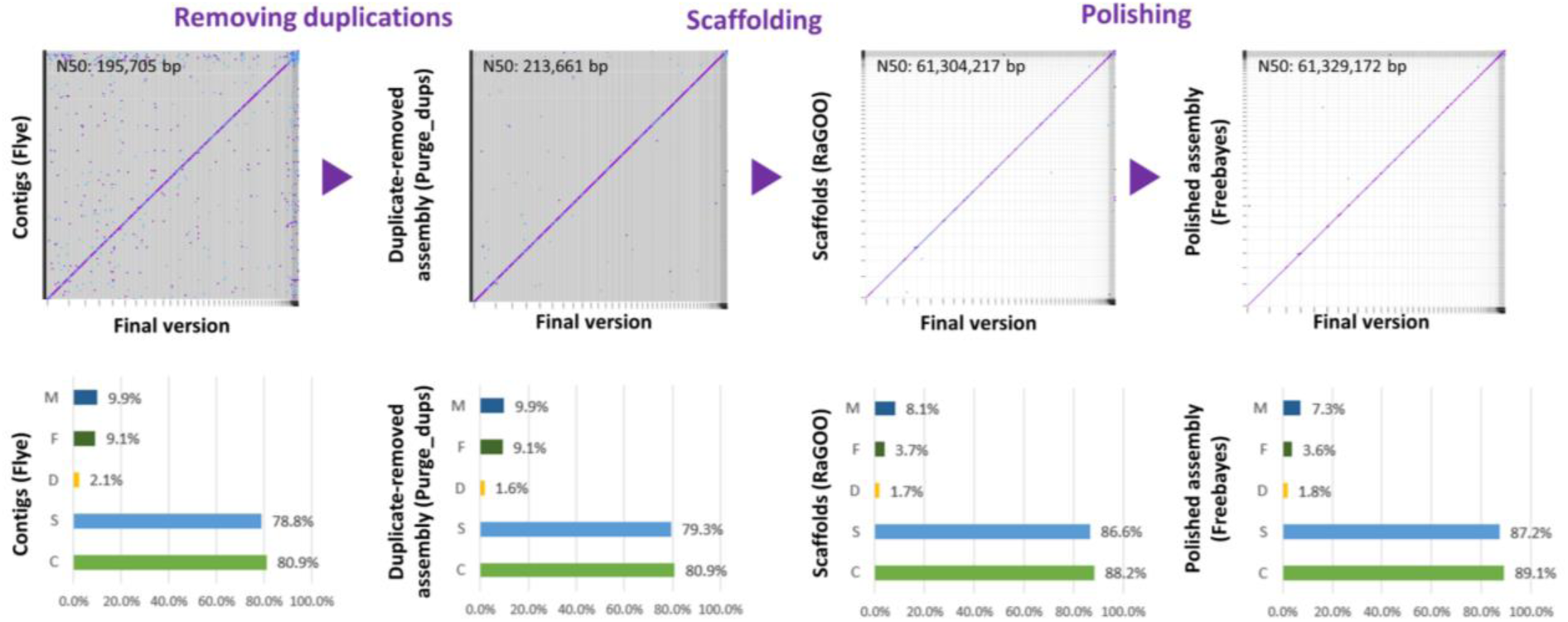
Genome assembly process. Improvement of the new genome assembly throughout the assembly process in terms of contiguity (contig N50), duplications and BUSCO gene contents.

**Figure S3.**
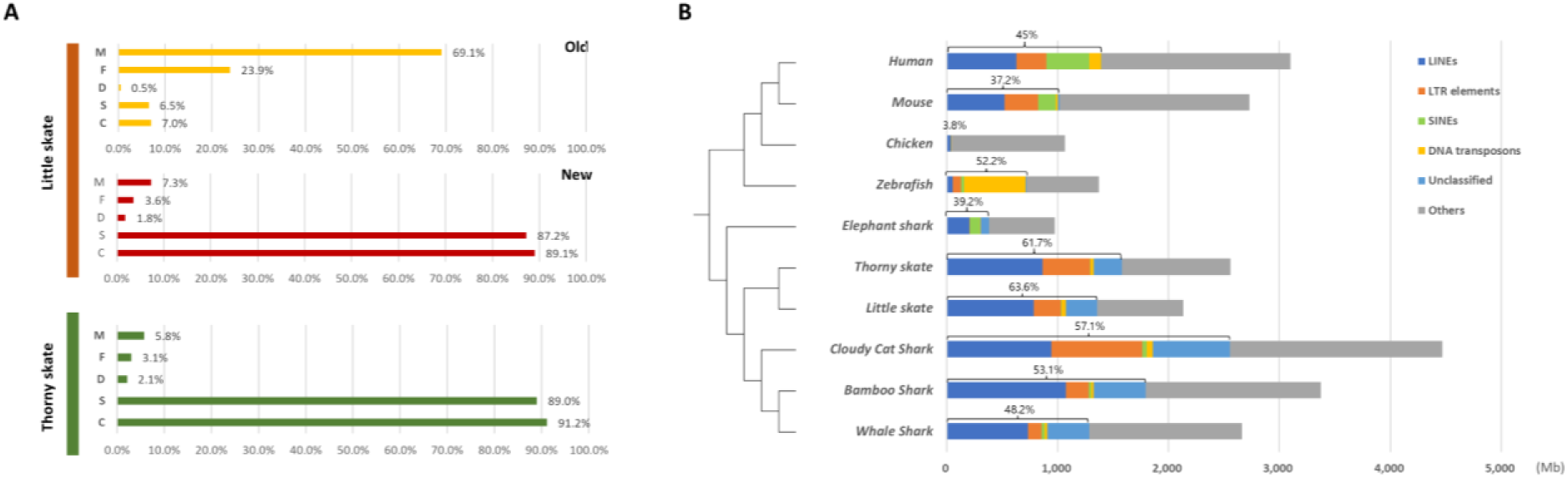
BUSCO genes and repetitive elements of little skate genome. A. The BUSCO gene contents summarizing the completeness of the new little skate genome. The y-axis represents different class of BUSCO genes as follows. M: Missing, F: Fragmented, D: Complete and duplicated, S: Complete and single-copy and C: Complete. B. Repetitive element identified in little skate compared with diverse phylogenetic lineage.

**Figure S4.**
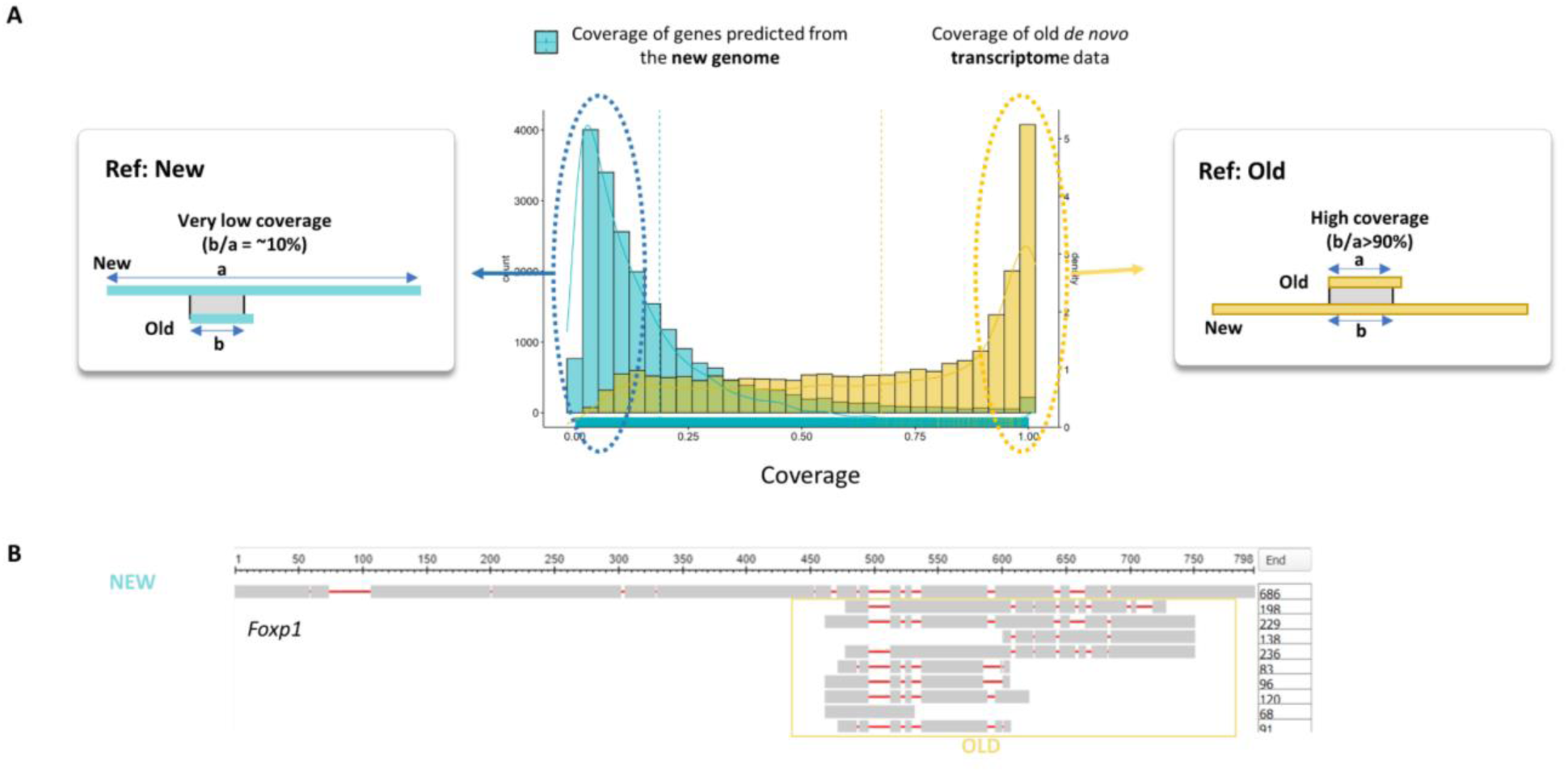
Limitation of previous transcriptome data. A. The Alignment coverage distribution when the alignment was performed using the predicted genes of new skate genome (blue) and old transcriptome (yellow) as the reference. B. The example case shown in *Foxp1*.

**Figure S5.**
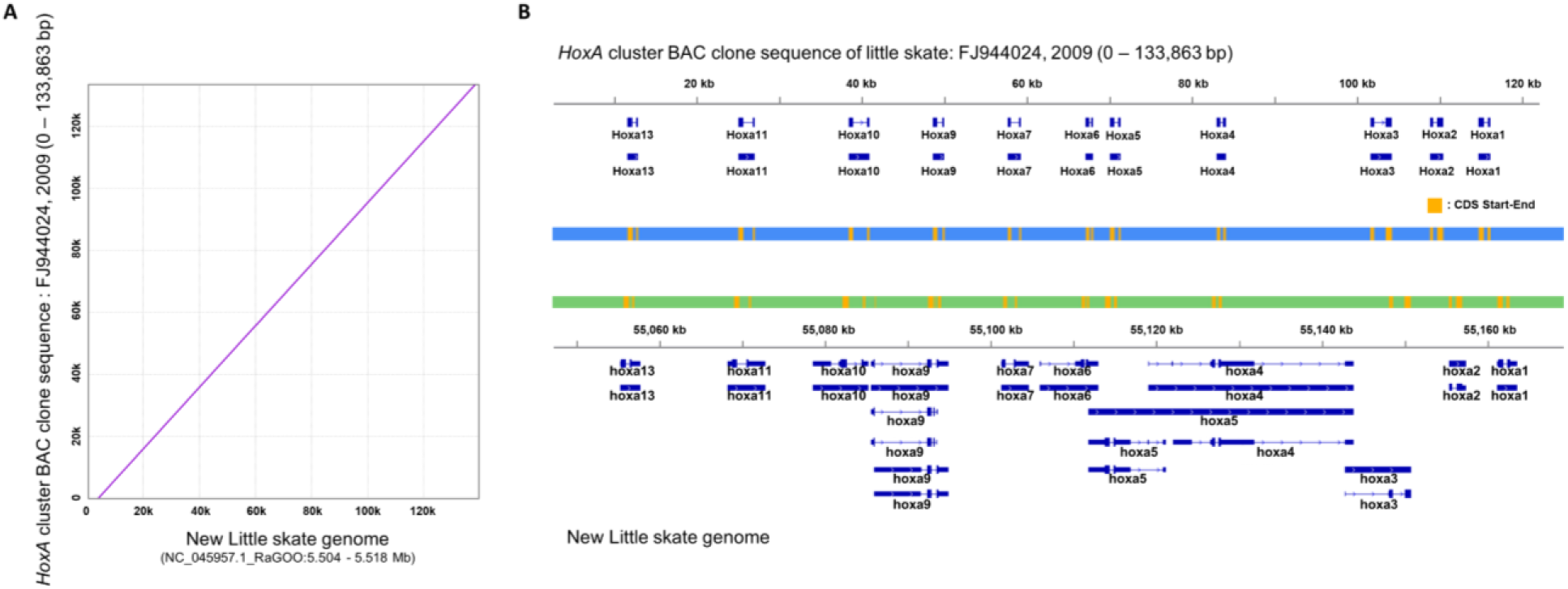
Qualitative assessment of genome assembly, *HoxA* cluster. A. Pairwise-alignment of *HoxA* cluster. B. *HoxA* genes of previous BAC clone and the new genome.

**Figure S6.**
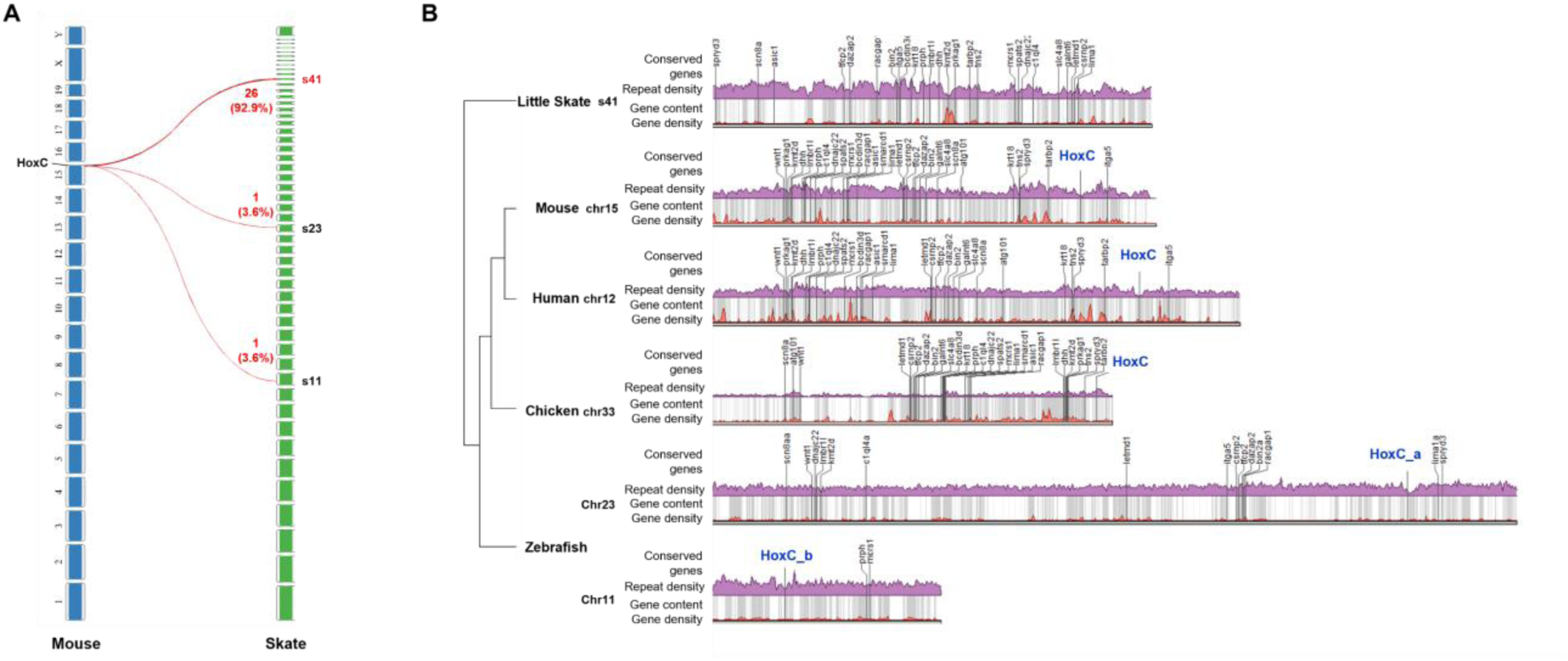
Comparison of *HoxC* clusters. A. The majority of ortholog genes near *HoxC* cluster highly conserved in multiple species are mapped to s41 of little skate genome. B. The repeat and gene contents near the orthologous genes. The *HoxC* cluster is in general located near the region with low repeat and gene contents except in chicken and zebrafish chr11.

**Figure S7.**
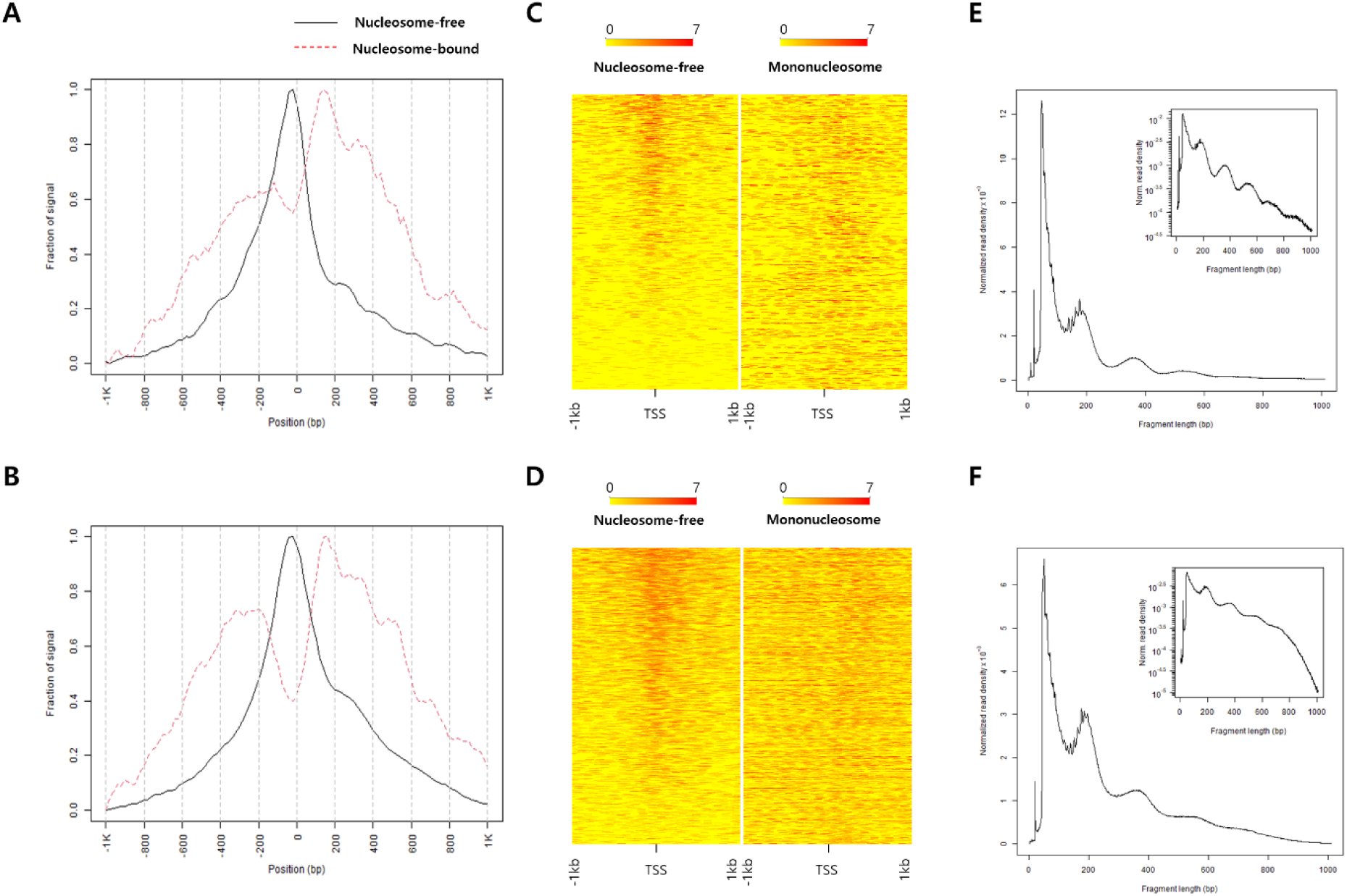
ATAC-seq data quality control. Fraction of signal plot showing ATAC-seq read depth nearby TSS sites of A. fin-MN and B. tail-SC. The heatmap of 1kb up and downstream regions of TSS sites in C. fin-MN and in D. tail-SC. Distribution of fragment size of the ATAC-seq data from E. fin-MN and F. tail-SC.

**Figure S8.**
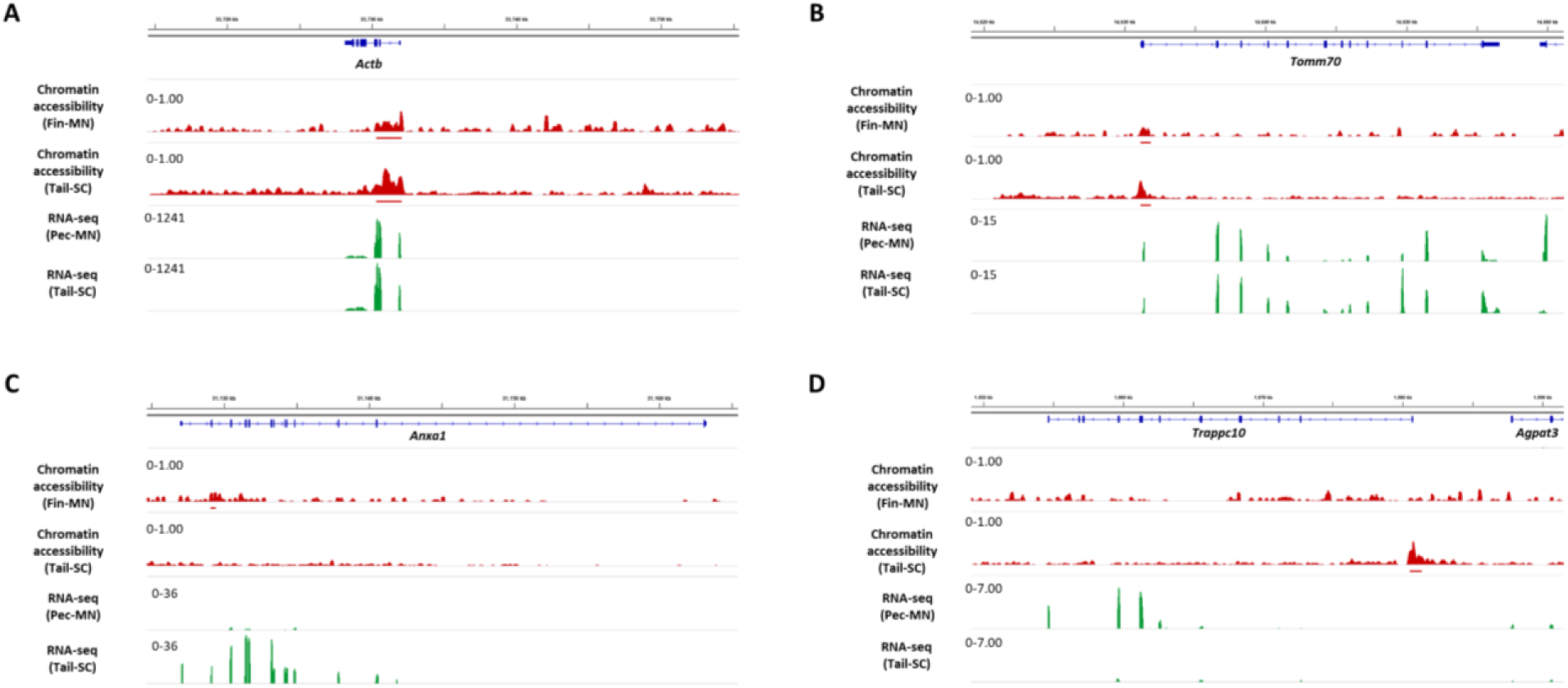
Chromatin accessibility of constitutively expressed genes and example genes with different peak types. The gene expression and chromatin accessibility of constitutively expressed genes (including shared ACRs), A. *Actb*, B. *Tomm70*, and example genes with fin-MN and tail-SC ACRs, C. *Anxa1,* D. *Trappc10*.

**Figure S9.**
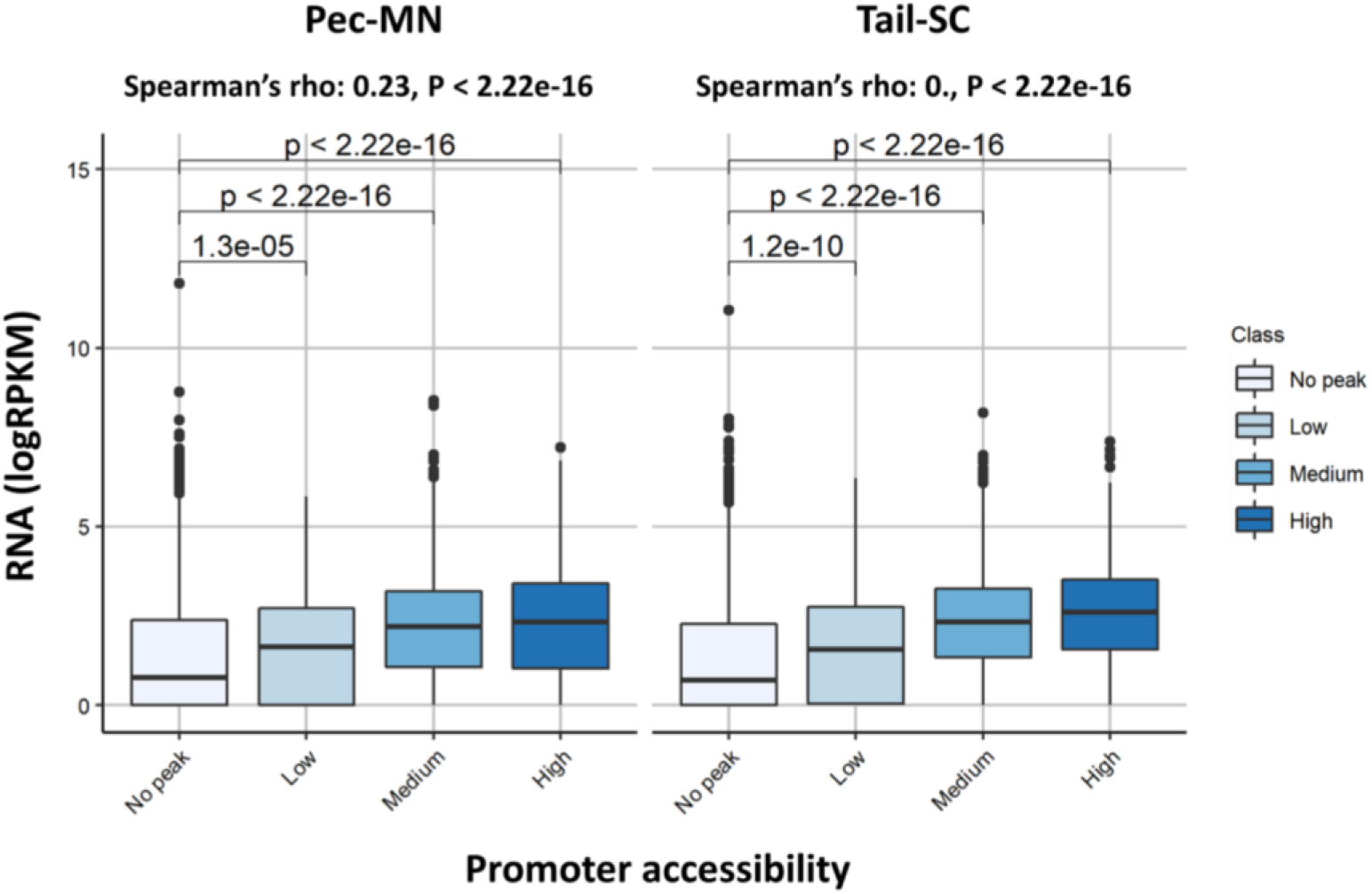
Correlation between promoter accessibility with gene expression. No peak indicates genes without any ACR located in promoters and low, medium and high are those genes with ATAC-seq read depth <Q10, Q10 <x< Q90 and Q90<. Center line represents median and the box limits shows upper and lower quartiles. The distance to maximum or minimum data point within the 1.5x interquartile range was used for whiskers.

**Figure S10.**
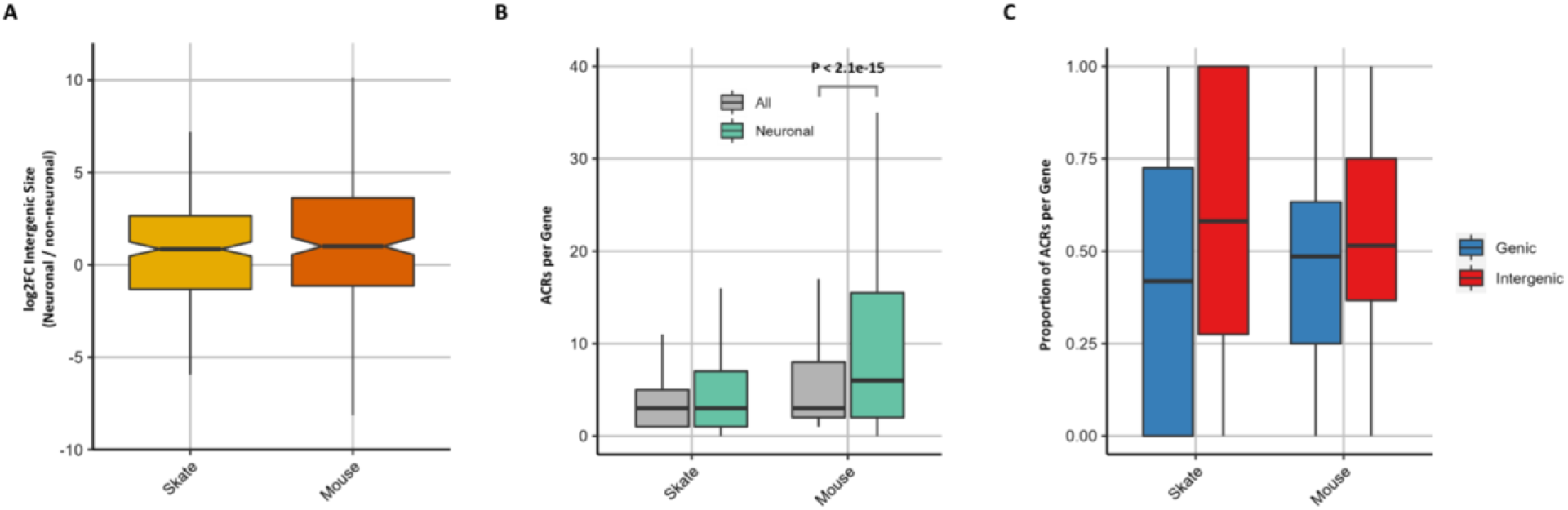
Comparison of intergenic size and number of ACRs between mouse and little skate. A. Comparison of intergenic size of orthologous genes of mouse and little skate. B. Distribution of the number of ACRs in all and neuronal genes. C. Proportion of genic and intergenic ACRs for each gene.

**Supplementary Table S1.**
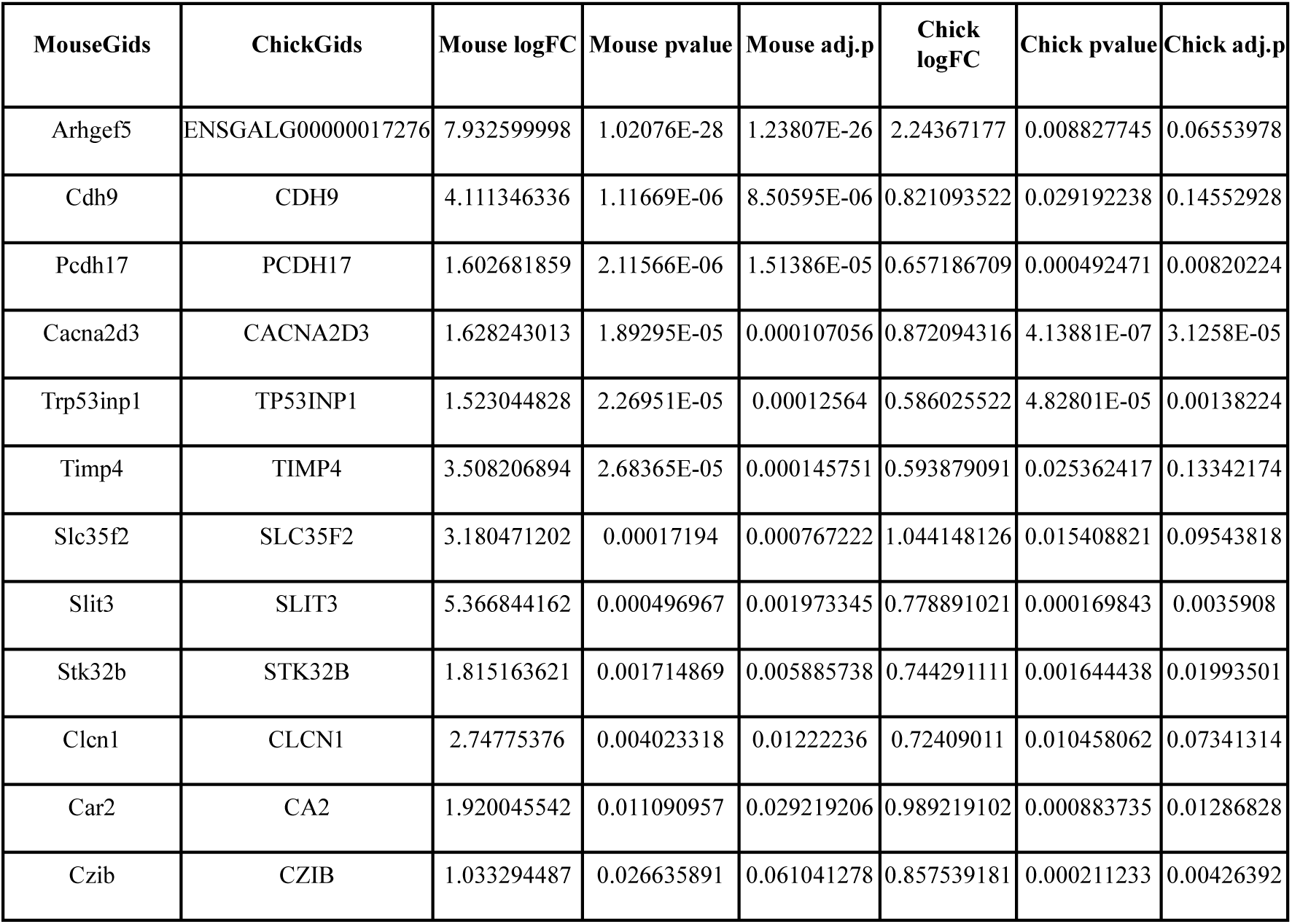
Motor neuron DEGs specifically lost in Skate.

**Supplementary Table S2.**
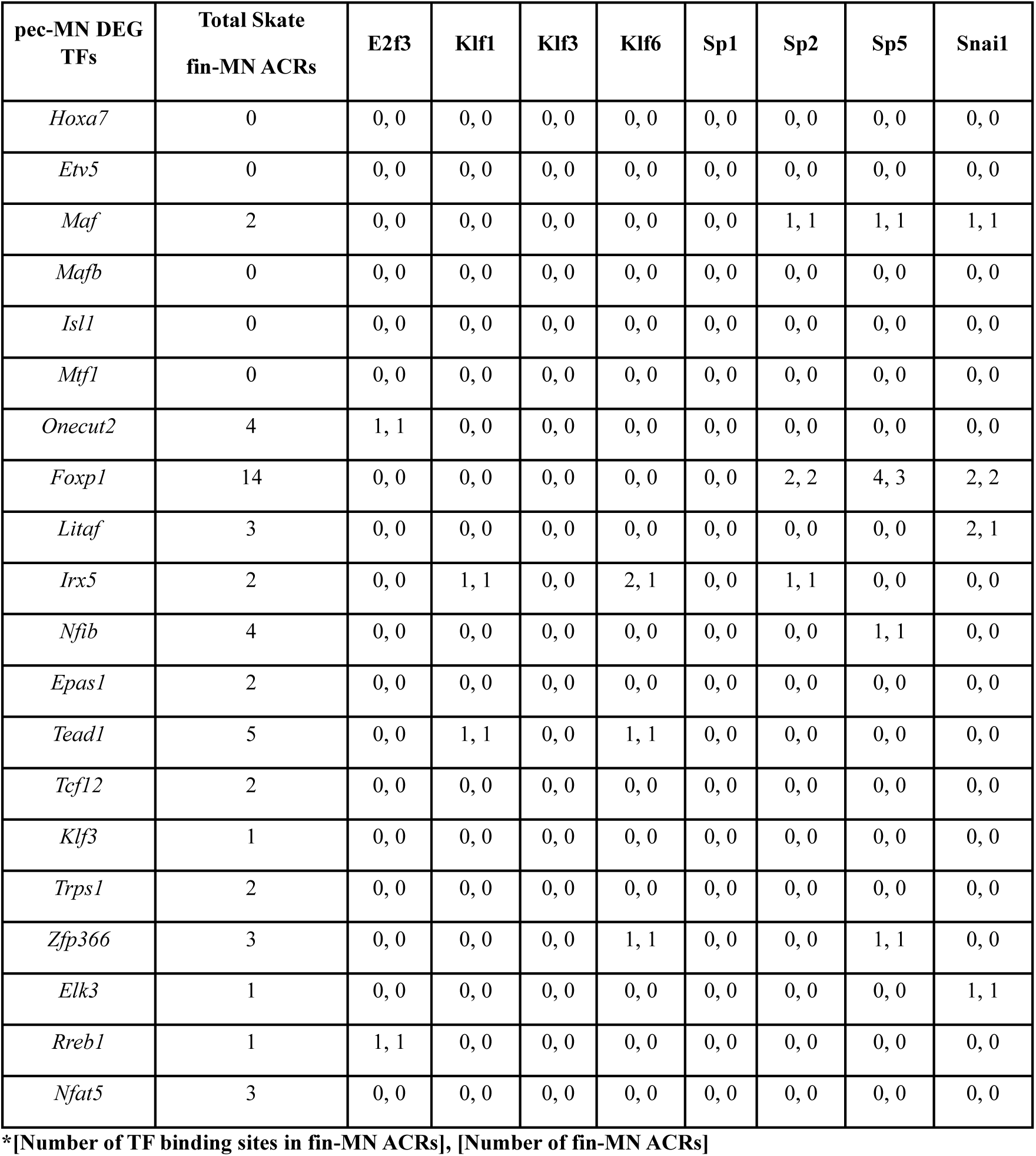
Putative binding site prediction in TFs of skate pec-MN DEGs.

**Supplementary Table S3.**
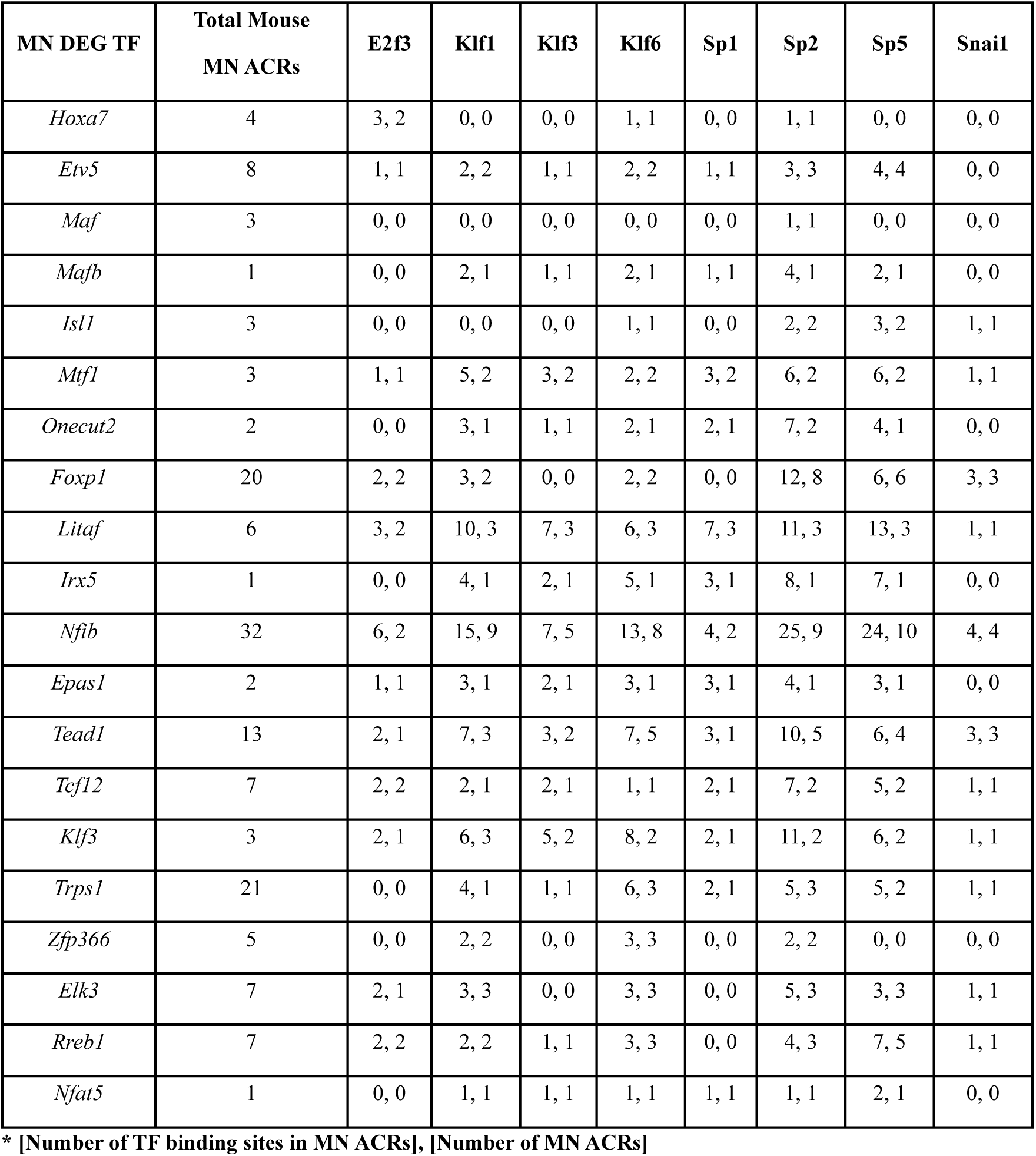
Putative binding site prediction in TFs of mouse MN DEGs.

## Notes

### Competing Interest Statement

The authors have declared no competing interest.

